# SOCIALITY, COMMUNITIES AND MORPHOLOGICAL FEATURES OF ANTS FROM MID-CRETACEOUS BURMESE AMBER

**DOI:** 10.1101/2022.02.10.479968

**Authors:** K.S. Perfilieva

**Author notes:** Translated into English by O.A. Belyaev (MSU, Dept. of Entomology).

## Abstract

We consider morphological diversity of ants from mid-Cretaceous Burmese amber. An eco-ethological hypothesis concerning its origin and features of Mesozoic and Cenozoic ant communities is proposed. It appears that some morphological features of representatives of the stem taxa allow us to speak about the absence of effective communication and, subsequently, group foraging in these ants. Therefore, the diversity of primitive Cretaceous ants, as predators, reflects their food specialization according to types of prey, on condition of their social lifestyle, that results in division of the ecological space among ant species into ecological niches. The occurrence of both effective communication and group foraging (mobilization) in the crown ant taxa, as crucial adaptation, has permitted them to exceed the bounds of niches of specialized predators, since type and size of prey are not strictly correlated to size of an ant and its mandibles; it also has given a chance to maintain large colonies. Due to this, myrmecocomplexes of modern ants are arranged on the principle of colonies dominance rather than the principle of division of ecological niches, like Mesozoic.

---

> *In memory of G.M. Dlussky, A.A. Zakharov, E.O. Wilson – the giants on whose shoulders we stand*

Burmese amber (Burmite, Kachin amber), ca. 99 Myr old, is rich with fossil organisms of amazing preservation and taxonomic diversity; it gives an insight into taxonomic diversity of orictocenosis, and what is more, provides a rare opportunity to study the structure of extinct biocenoses. Now the described animals comprise 651 families, 1382 genera, 2038 species; among them arthropods amount to 583, 1264 and 1908, accordingly (Ross, 2021). Ants are represented in Burmite by one family Formicidae: 31 species of three extinct subfamilies (Haidomyrmecinae, Zigrasimeciinae, Sphecomyrminae) and moreover, judging by a certain unpublished data, at least three crown subfamilies (Ponerinae, Dolichoderinae, Formicinae) (although the data on crown subfamilies is supposedly true only for late Burmese amber – Tilin amber, 72 Myr) (Zhang et al., 2018; Zheng et al., 2018; Boundinot et al., 2020) (Tab.). Recently two genera (*Camelomecia, Camelosphecia*) were also discovered; they were described on winged sexual individuals (two females and one male of three species) referred to the superfamily Formicoidea, in the quality of a sister group to Formicidae (Barden and Grimaldi, 2016; Boudinot et al., 2020). I have doubts on referring insects without metapleural glands, such as *Camelomecia* and *Camelosphecia*, to formicoids, besides non-formicoid venation of wings in *Camelomecia* also should be noted. Taxonomic and morphological diversity (unique morphology and variety of mandibles, in particular) of formicoids in Burmite has no satisfactory explanation to the moment, regardless of the fact that questions of how and why such morphological diversity occurred are raised in almost every publication containing descriptions of new ant species from Burmese amber, since modern ants, although taxonomically rich (about 14,000 recent species, 17 subfamilies), have no analogues of such mandibles. The mandibles of all modern ants have common features; even taking into account the specialized forms, modifications of the mandibles are quite well-studied and organized into morphological series with common root (Dlussky and Fedoseeva, 1988). The purpose of this study was the analysis of representatives of the Burmese amber myrmecofauna along with the causes of specific morphological radiation of the stem taxa and their extinction.

**Table.**
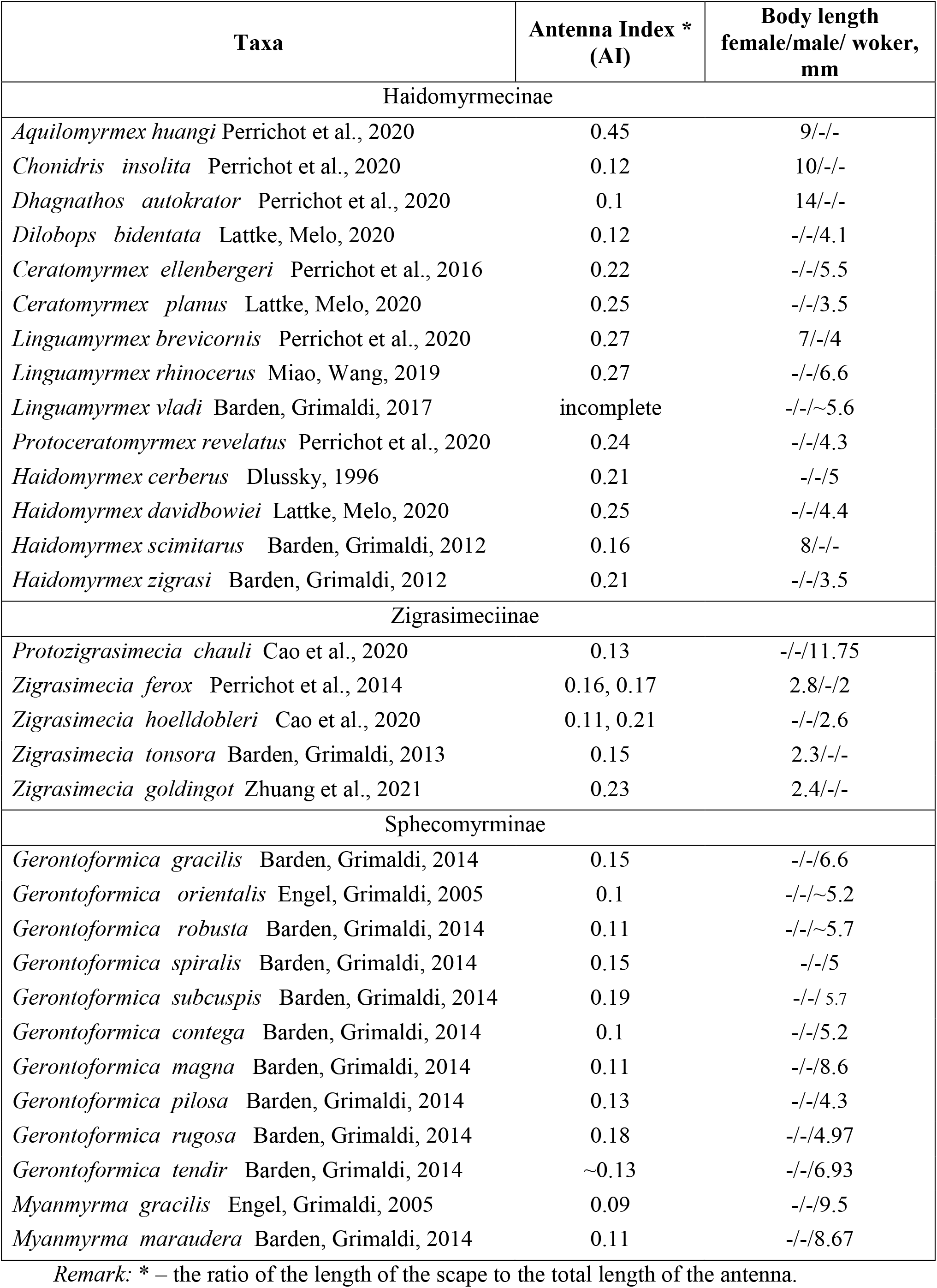
Taxa and some morphometric characteristic of Burmite formicoids.

| Taxa | Antenna Index *<br>(AI) | Body length<br>female/male/ worker,<br>mm |
| --- | --- | --- |
| Haidomyrmecinae |  |  |
| *Aquilomyrmex huangi* Perrichot et al., 2020 | 0.45 | 9/-/- |
| *Chonidris insolita* Perrichot et al., 2020 | 0.12 | 10/-/- |
| *Dhagnathos autokrator* Perrichot et al., 2020 | 0.1 | 14/-/- |
| *Dilobops bidentata* Lattke, Melo, 2020 | 0.12 | -/-/4.1 |
| *Ceratomyrmex ellenbergeri* Perrichot et al., 2016 | 0.22 | -/-/5.5 |
| *Ceratomyrmex planus* Lattke, Melo, 2020 | 0.25 | -/-/3.5 |
| *Linguamyrmex brevicornis* Perrichot et al., 2020 | 0.27 | 7/-/4 |
| *Linguamyrmex rhinocerus* Miao, Wang, 2019 | 0.27 | -/-/6.6 |
| *Linguamyrmex vladi* Barden, Grimaldi, 2017 | incomplete | -/-/~5.6 |
| *Protoceratomyrmex revelatus* Perrichot et al., 2020 | 0.24 | -/-/4.3 |
| *Haidomyrmex cerberus* Dlussky, 1996 | 0.21 | -/-/5 |
| *Haidomyrmex davidbowiei* Lattke, Melo, 2020 | 0.25 | -/-/4.4 |
| *Haidomyrmex scimitarus* Barden, Grimaldi, 2012 | 0.16 | 8/-/- |
| *Haidomyrmex zigrasi* Barden, Grimaldi, 2012 | 0.21 | -/-/3.5 |
| Zigrasimeciinae |  |  |
| *Protozigrasimecia chauli* Cao et al., 2020 | 0.13 | -/-/11.75 |
| *Zigrasimecia ferox* Perrichot et al., 2014 | 0.16, 0.17 | 2.8/-/2 |
| *Zigrasimecia hoelldobleri* Cao et al., 2020 | 0.11, 0.21 | -/-/2.6 |
| *Zigrasimecia tonsora* Barden, Grimaldi, 2013 | 0.15 | 2.3/-/- |
| *Zigrasimecia goldingot* Zhuang et al., 2021 | 0.23 | 2.4/-/- |
| Sphecomyrminae |  |  |
| *Gerontoformica gracilis* Barden, Grimaldi, 2014 | 0.15 | -/-/6.6 |
| *Gerontoformica orientalis* Engel, Grimaldi, 2005 | 0.1 | -/-/~5.2 |
| *Gerontoformica robusta* Barden, Grimaldi, 2014 | 0.11 | -/-/~5.7 |
| *Gerontoformica spiralis* Barden, Grimaldi, 2014 | 0.15 | -/-/5 |
| *Gerontoformica subcuspis* Barden, Grimaldi, 2014 | 0.19 | -/-/ 5.7 |
| *Gerontoformica contega* Barden, Grimaldi, 2014 | 0.1 | -/-/5.2 |
| *Gerontoformica magna* Barden, Grimaldi, 2014 | 0.11 | -/-/8.6 |
| *Gerontoformica pilosa* Barden, Grimaldi, 2014 | 0.13 | -/-/4.3 |
| *Gerontoformica rugosa* Barden, Grimaldi, 2014 | 0.18 | -/-/4.97 |
| *Gerontoformica tendir* Barden, Grimaldi, 2014 | ~0.13 | -/-/6.93 |
| *Myanmyrma gracilis* Engel, Grimaldi, 2005 | 0.09 | -/-/9.5 |
| *Myanmyrma maraudera* Barden, Grimaldi, 2014 | 0.11 | -/-/8.67 |

Remark: \* – the ratio of the length of the scape to the total length of the antenna.

## Material and methods

I have analyzed published materials of Cretaceous and modern ants. Images for drawing were taken from the AntWeb (https://www.antweb.org) with indication numbers of specimens, also from our archive. Drawings and measurements of morphological structures were performed using the Inkscape program distributed under a free license. Graphs were created in Excel 2013. Measurements of structures and obtainance of missing information were based on the published images as well as on papers of various authors about descriptions of the corresponding Cretaceous ants (see Tab.). I have taken the sample of modern ants for measurements from the article about the comparison of morphospaces of stem and modern ants (Barden at al., 2020).

## Results and discussion

### Cretaceous ant diversity

The Cretaceous period is characterized by significant changes in entomofauna that appeared from a change in the dominant groups of plants in phytocenoses, followed by a change in the structure of biocenoses. The proportion of modern families in the entomofaunas since the Early Cretaceous would increase from half to three fourths, it points at the formation of a modern appearance of entomofauna at the family level in this period (Zherikhin, 2003). The increase in diversity of phyllophagous insects, parasitoids and other hymenopterans is observed for the Cretaceous period. Burmese amber shows an significant diversity of arthropods from the Middle Cretaceous (Ross, 2021). Calculations of arthropod biodiversity in Burmite at the family level indicate high values for the occurrence of new families and “fauna turnover” (the sum of families first time encountered and last time referred to the total number of families in a given locality) in this amber (Rasnitsyn, 2016). These facts point at a diverse and rich resource base for terrestrial predatory insects at this period.

Cretaceous ants are represented not only by the above listed extinct subfamilies, but also by several crown subfamilies. Ant specimens from extinct genera, still not referred to any subfamily, are described, too, although this is rather the result of a technical character – authors of the latest phylogenetic schemes have not studied yet these individuals for the presence of features they have emphasized (specimens from the Taimyr amber, Armaniinae). There are described species from the Cretaceous crown groups: two species of Ponerinae (Dlussky, 1999), and by one species of Dolichoderinae (McKeller et al., 2013), Formicinae (Grimaldi and Agosti, 2000), Aneuretinae (Engel and Grimaldi, 2005), and Myrmicinae (Dlussky, et al., 2004). The Cretaceous ant fauna also revealed intermediate forms – Armaniinae (Formicidae in the current system) together with recently described winged representatives of the clade which includes two genera (*Camelomecia* and *Camelosphecia*) and is sister to Formicidae. So, the myrmecofauna of the Cretaceous already included representatives of the main crown subfamilies, besides, the major taxonomic and morphological diversity was demonstrated by the extinct subfamilies with a little over 50 described species to the moment. On this basis it can be expected that the Cretaceous was the time of “formicoidization” (similar to evolutionary phenomena of “arthropodization”, “ornitization” etc.), i.e. the period of occurrence of certain features specific to modern ants in unusual combinations in different phylogenetic branches; the entire set of such features was formed in crown ant taxa, which are considered a monophyletic group.

The proportion of ants in orictocenoses would grow throughout the entire period of existence of the family from the end of the Lower Cretaceous to the present time, but in the Cretaceous most commonly it equaled first fractions of a percent, numerically several specimens in general (LaPoll and Dlussky, 2013). Our preliminary calculations show that ants make up 2.6% of all arthropod species in Burmese (Kachin) amber. According to studies of hymenopterans in Burmese amber, ants make up about 9.1% of all hymenopteran specimens (Zhang Q. et al. 2018). The important thing is that parasitoids constitute larger or comparable proportion of specimens among hymenopterans (Scelionidae, 16.1%; Chrysididae, 11.5%; Bethylidae, 7%) in view of evaluation of resource base richness of a biocenosis. Therefore, the proportion of ants in the taxonomic diversity of Burmite along with the number of individuals were quite noticeable against the background of the general arthropod diversity, although not comparable with the current state of this group of insects (for example, the proportion of ant specimens in Dominican amber is about 36%, while the biomass of ants in a present-day tropical forest constitute 15-20% of all animals) (Hölldobler and Wilson, 1990; LaPolla and Dlussky, 2013).

The finding of *Aethiocarenus burmanicus* Poinar, Brown 2017 serves a clear evidence of significant ecological role of ants in Burmite paleobiocenoses, since it demonstrates myrmecomorphy, which is common among modern harmless arthropods for self-defense purposes (a case of Batesian mimicry) (Vršanský et al., 2018). Still not an ant from crown groups was described from Burmese (Kachin) amber, although they were found in later Burmese amber of Tilin (Zheng D. et al., 2018).

### Ant origin hypotheses

The best-known and widely accepted hypothesis on the conditions of occurrence of ants was proposed by Wilson and Hölldobler (Wilson and Hölldobler, 2005), it is known as the “Dynastic-Succession Hypothesis” (DSH). According to DSH, the main ancestral group of modern ants is represented by the forms that have favored ground and leaf-litter sites for predatory lifestyle (geobionts, stratobionts), specifically ponerines^1^, which have arisen in the Middle Cretaceous, spread over the world in the Paleogene and eventually given rise to modern subfamilies, followed by a transition to the ecological dominance of ants along with a diet change due to the expansion of angiosperms in tropical regions. There are two reasons for this conclusion: the litter is a habitat with very large biomass, where arthropod predators, such as Cretaceous ants and present-day primitive taxa (specifically poneromorph ants), are capable to feed themselves. The second reason, called “The Ponerine Paradox” by the authors, consists in the contradiction (as the authors considered) between the fact of the wide geographical distribution of ponerine taxa with regard of their primitive social organization: small-populated monogynous colonies, absence of polymorphism and polyethism, solitary hunting, primitive communication – alarm signal, moreover trophallaxis and mobilization (recruitment) of workers to a food source are almost absent. This implies that the origin of modern ants from primitive “ponerines” adapted to hunting in the litter after the radiation of this group in the Late Cretaceous and the Paleocene could explain the wide geographical distribution of the primitive group of poneromorphs as well as the holding of positions in competition with later spread (occurred?) more progressive myrmicines, dolichoderines, and formicines, which have appeared due to a diet change linked to a distribution of honeydew-producing insects, and could not completely displace the well-adapted poneromorphs, which shared ecological niches in their habitat.

The position concerning the origin of modern ants from the specialized leaf-litter predators was not supported by Dlussky (Dlussky and Rasnitsyn, 2007). He pointed out that the Cretaceous ants found by that time, sphecomyrmines, possessed the habitus of a terrestrial predator, successfully foraging on litter surface and in tree layer, i.e. were not similar to litter dwellers. He suggested the formation of modern main phylogenetic branches as a life forms adapted to hunting in different strata of a biocenosis: in the soil («poneromorphs»), in the leaf-litter (aneuretines), on the soil surface and in the trees (formicines, dolichoderines, and myrmicines) (Dlussky and Fedoseeva, 1988).

It is obvious now that paleontology does not support Wilson’s assumption of earlier origin of poneromorph taxa. Also no predominance of poneromorph representatives is evidenced among Cretaceous ants, while representatives of the “progressive and younger” (in the framework of the hypothesis of Wilson and Hölldobler) subfamilies are presented in Cretaceous orictocenoses, with well-defined features indicating their belonging to crown subfamilies – Formicinae, Dolichoderinae, Myrmicinae, Aneuretinae. Clearly that the resulted absence of litter-dweller forms in the fossil record is quite consistent, it cannot be the main argument. However, the assertion about formation of crown subfamilies of ants due to adaptations to predation in the litter seems rather controversial. As can be seen from modern taxa, the morphology of ants living and hunting in the litter changes towards decrease in the length of antennae, decrease in the relative eye size, thickening of the integument, specialization of the mandibules. It is not like the features that all crown groups of ants have inherited. However, I do not deny the connection of ancestral taxa of ants with the leaf-litter, I only draw attention to the absence of morphological specialization in ancestral group for life in the litter. A somewhat different picture emerges against the described one in the hypothesis of dynastic succession. The ancestors of modern crown groups should have inherited morphological features that for some reason remained fixed in all (almost without exceptions) modern crown groups: three(and more)-toothed mandibles, geniculate antennae with an elongated scape, the structure of the funicullus (relatively long curved pedicel, enlarged apical segments), the presence of ocelli and compound eyes. In modern ants, with several changed features or one only, it is the result of specialization and modification (Dlussky, Fedoseeva, 1988). The hypothesis of the origin of ants should explain their presence. It appears that this complex of features is unnecessary in the litter environment and within primitive colonies, like in some «poneromorphs», consequently it could not be formed.

### Morphological features of ants and relationship with sociality. Hypothesis on the origin of crown ant groups and their evolutionary success

Some authors, although recognizing the DSH of Wilson and Hölldobler as a working assumption, contradict it in a paradoxical way. Thus, Perrichot with co-authors, as one of the leading professional researchers of Cretaceous formicoids, being the authors of stem formicoids ecology and morphology analyses, on the one hand accept the DSH. On the other hand, they quite fairly assume that ants within Formicoidea developed like a form with *“innovation suite for cursorial or surface-based predation”* (Boudinot et al., 2020). Therefore, Cretaceous formicoids have a certain set of morphological features, including prognathy (this feature is considered in the paper of Fedoseeva E. (2001)), rotation of the antennal toruli laterad (however, not all Burmite ants possessed this modality, Fig. 1*a*, *d, h*), elongation of the procoxae and other. The authors relied on cladistic analysis (i.e. revealed synapomorphies for the entire group), apparently for this reason they did not include morphological features, which are present in other Aculeata, too, but also characterize a set of features of fast cursorial surface-based predators: for example, large compound eyes and ocelli, long antennae and legs. Now 31 ant species are currently described from the Burmese amber (Kachin amber) (see Tab.). I do not include the three species of the genera *Camelomecia* and *Camelosphecia* into the analysis due to the absence of metapleural glands, the non-formicoid wing venation in *Camelomecia*, the lack of evidence for the presence of a wingless caste, and, consequently, doubts about the necessity of referring them to Formicoidea.

**Fig. 1.**
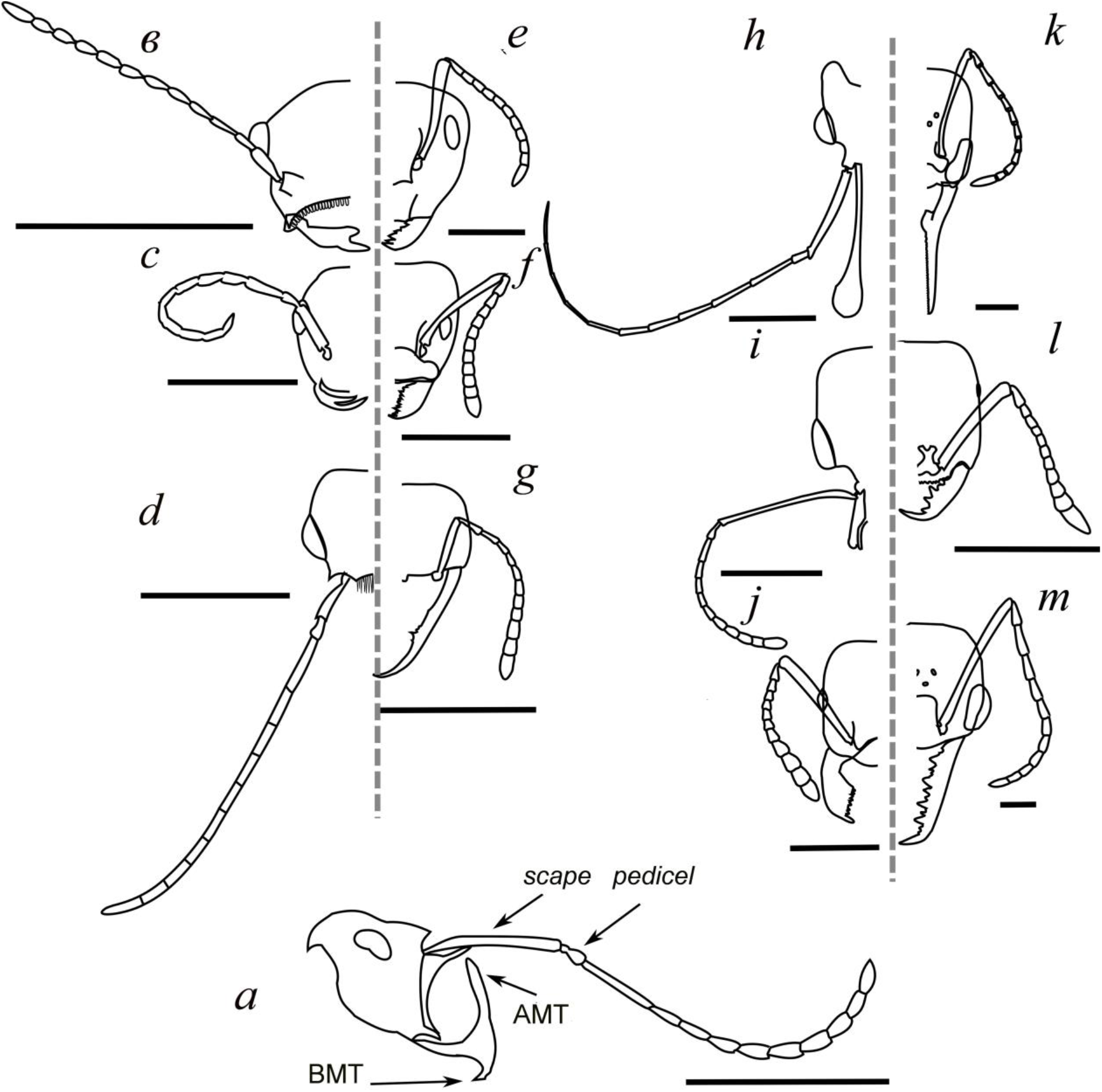
Head drawings from profile (*a*) and dorsal (*b-m*) views of Cretaceous (*a-d, h, i*) and modern (*e-g*, *j-m*) ants based on photo images. Morphology of antennae, mandibules, clypeus, form and size of eyes and ocelli is displayed: *a* – *Linguamyrmex brevicornis* (FANTWEB00035); *b* – *Zigrasimecia tonsora* (ANTWEB1008098); *c* – *Gerontoformica orientalis* (JWJ-BU19); *d* – *Dilobops bidentata* (FANTWEB00039); *e* – *Camponotus abrahami* (CASENT0910439); *f* – *Azteca adrepens* (CASENT0173823); *g* – *Martialis heureka* (CASENT0106181); *h* – *Ceratomyrmex ellenbergeri* (NIGP164022); *i* – *Aquilomyrmex huangi* (FANTWEB00024); *j* – *Manica bradleyi* (CASENT0106022); *k* – *Harpegnathos saltator* (CASENT0101783); *l* – *Onychomyrmex* sp. (CASENT0069959); *m* – *Myrmecia* sp. (CASENT0006136). Scale bar: 1 mm. Abbreviations: AMT, BMT – apical and basal mandibular tooth; AntWeb database numbers of specimens given in brackets.

All researchers of the Burmite myrmecofauna are consentaneous in assessment of the ecological role of the described ants. The variety of sizes of ants and their mandibles indicates specialized predation. The sociality of Cretaceous formicoids has been discussed since the first descriptions (Dlussky, 1984, 1987; Wilson et al., 1967; Wilson, 1985). Barden and Grimaldi summarized in their study the morphological evidence for eusociality in Cretaceous formicoids (at least some of them are known for having winged and wingless castes, besides the reproductive females are notable for traces of discarded wings), also they suggested that syninclusions of rather rare ants in Cretaceous orictocenoses in one piece of resin indirectly evidence of sociality and group behavior (Barden and Grimaldi, 2016). The mentioned morphological characteristics certainly point at the sociality of Cretaceous formicoids, although the assumption of group behavior seems baseless. Four Burmite pieces are discussed in the article: JZC Bu1814 – 6 individuals of *Gerontoformica spiralis;* JZC Bu116 – 11 individuals of *G. spiralis* and 1 worker *Haidomyrmex zigrasi*; JZC Bu1645 – 21 individuals of *G. orientalis*, *G. contegus*, *G robustus*; JZC Bu1646 – two fighting workers of *G. tendir* and *G. spiralis*. The mutual presence of several individuals of rare (according to the authors, but in the light of the facts presented above, this is not so) for the biocenoses ants of the same species together with the two fighting ants are interpreted by researchers as coordination of actions during foraging and aggressive interactions between species respectively; however, the authors come to the conclusion that a pheromone trail apparently was not applied. It should be noted that the authors assume that the crown taxa ecologically have replaced the stem taxa, though they do not describe the mechanisms of replacement, except for the remark about the ants with specialized mouthparts (Zigrasimeciinae, Haidomyrmecinae) which obviously depended on food sources. It appears to me that the presence of many (scores) other animals (among them a snail and arthropods such as a spider, a cockroach, a scolebythid wasp, beetles, springtails etc.) in the considered small pieces of amber makes the second assumption of the authors about the aggregation close to the food source more convincing. The aggregation of ants of two or three species from neighboring nests considering their social lifestyle does not appear to be an astronomically extraordinary event even in the case of absence of mobilization. The second peculiarity, the presence of worker ants of different species in one piece without signs of aggression, also supports this assumption and excludes the protection of a food source. Therefore the presence of eusociality seems to be entirely proven, however, the protection of food sources does not constitute a regular attribute of coexistence in the described ant communities (equally as the mobilization to a food source). Apparently, protection of food sources exhibits at the level of certain individuals. Indeed, the syninclusions of two ants badly injured in the fight reveal us that the struggle lasted for quite a long time, even so these ants remained one on one, while modern ants in such situations most commonly have the assistance, that was also reflected in the Eocene amber (for example, Radchenko and Perkovsky, 2021). Obviously primitive polymorphism based on isometric size variation of workers could also be observed among stem ant taxa (Cao et al., 2020b).

The available data on modern ants allowed to distinguish several morphological features and trends that characterize crown groups of ants, which represent the majority of modern species and dominate in all biocenoses (Myrmicinae, Dolichoderinae, Formicinae):

1. Lateral turn of antennae, geniculate antennae with elongated scape (AI>3). (apomorphy – present in all crown ant taxa, though in stem ant taxa the morphology of antennae is diverse, usually antennae filiform, very long, scape may be extra short AI 0.1– 0.25 (see Tab., Fig. 1)).
2. Morphology of funiculus – relatively long curved pedicel and increase in size of apical segments of flagellum (synapomorphy?) (see Fig. 1).
3. Relatively small mandibles with at least three teeth (synapomorphy?) (see Fig. 1).
4. Compound eyes reduced in size, besides ocelli present (tendency) (Fig. 2).
5. Trophallaxis (tendency in dominant subfamilies).
6. Licking and carrying larvae and pupae in the process of brood care (synapomorphy?)
7. Poisonous sting replaced with acid gland (tendency in dominant subfamilies).

**Fig. 2.**
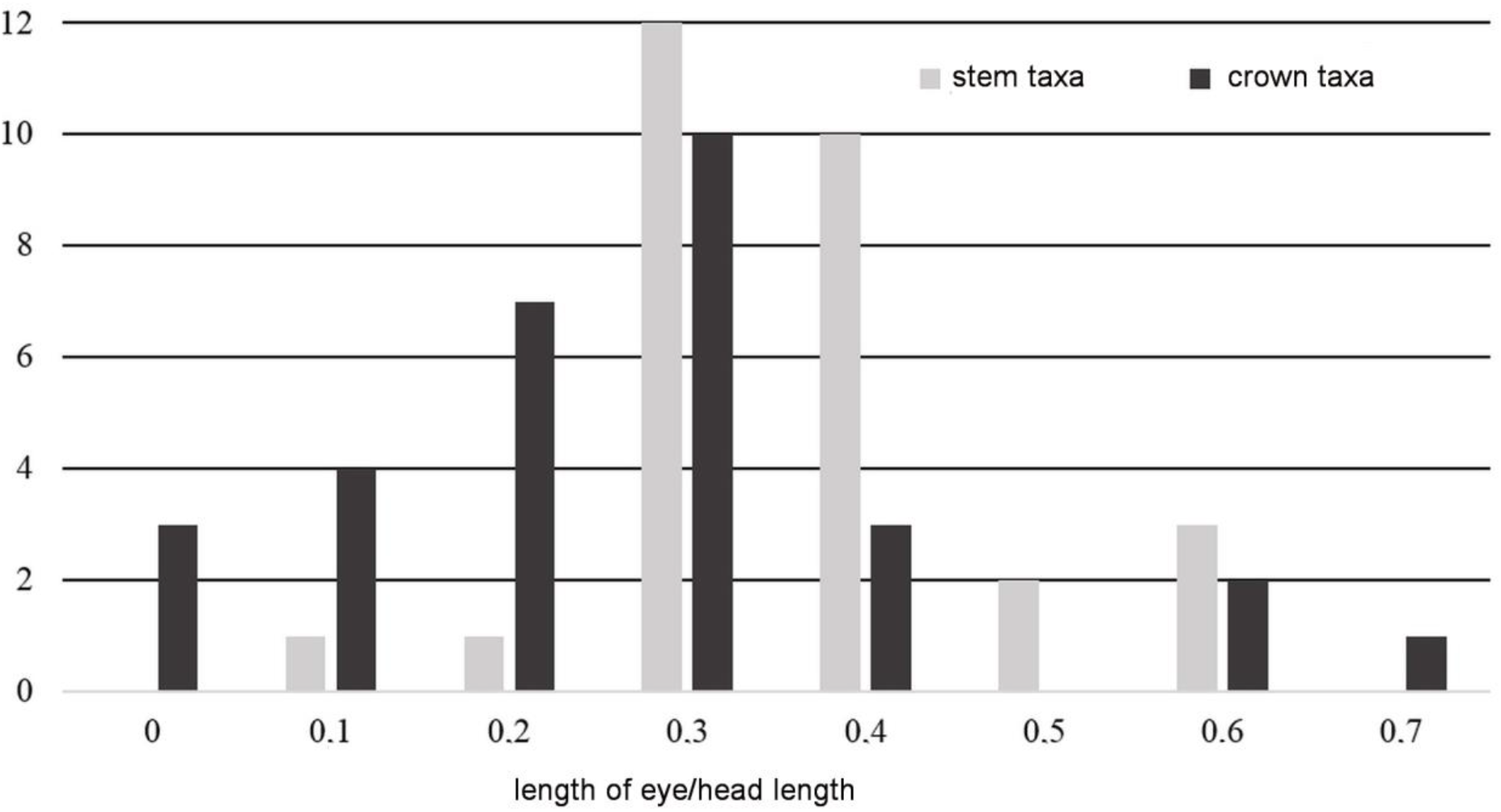
Eye length distribution in relation to head size in recent and Cretaceous formicoids. Scale bar: 1 mm.

Although sociality among ants is presented by a rather narrow range of variations, unlike other higher hymenopterans exhibiting all stages of the development of eusociality that can be traced from solitary way of life to giant-number bee colonies, still the features of a primitive social organization are observed in quite a big number of ant taxa. For example, the number of species of (primitive) poneromorphs in forest ecosystems constitute 22.2% (while the number of individuals is only 12.4%) (Ward, 2000). The most primitive ants are distinguished by the following features (the progressive taxa possess another modalities of these features): the presence of at least two castes of females, besides workers in such species quite often capable to change their status to that of reproductive females, they do not build a complex nest, get food by solitary predation, do not protect their foraging area, mobilization and trophallaxis absent; yet they recognize a common alarm signal, care of their brood, carry larvae and pupae, contact with each other upon meeting, perform grooming. And generally these species are indeed associated with the leaf-litter, in total it appears to confirm the DSH. However, all this does not explain the presence of general morphological features in ants as well as the tendencies listed above.

The concept proposed here is reduced to the idea of development of crown ants as represented by efficient fast cursorial surface-based wingless predators with more advanced social communication against the stem taxa. Dlussky suggested that geniculate antennae are associated with eusociality due to the possibility of manipulating small objects, it is hard to argue with this (Dlussky, 1984; Dlussky and Fedoseeva, 1988). They have shown that social wasps and bees, which care of their brood, have a relatively elongated scape (AI~0.3). However, ants for some reason required further increase in the length of scape (AI≥0.3, in more advanced dominant subfamilies Formicinae, Myrmicinae, and Dolichoderinae AI≥0.4) and in the lateral turn of antennae, while bees and wasps quite successfully coped with the tasks of manipulating small objects, although geniculate antennae are also observed in (nonsocial) parasitoid wasps, for example *Anastatus* sp. (Eupelmidae) exploiting insect eggs. Furthermore, termites being eusocial non-hymenopteran insects do not tend to develop geniculate antennae. Therefore, probably some other explanation is required concerning geniculate antennae and lateral turn of antennae in crown ant taxa. I suppose that a further increase in the relative length of scape and in the turn of antennae could provide better orientation for a fast cursorial surface-based wingless predator. It is possible that Johnston’s organ located in the pedicel could be one of the reason; it is a multisensory organizer, partially adopted functions of orientation from the visual organ. Johnston’s organ in ants responses for gravity perception as well as performs the functions of a wind compass and a step integrator (estimation of the passed distance) (Grob et al., 2020). From an engineering point of view, more convenient to analyze data from these receptors – in case when they farther located from each other. The compromise between both necessity to dispose the analyzers as far apart as possible and yet totally control the space directly near mandibles and mouth opening, has led to 1) a lateral turn of antennae, 2) an elongation of the scape, and 3) a change in shape of the pedicel – the elongated segment curved at the base allows funiculus to maximally approach scape, and consequently mandibles and mouth opening.

The third peculiarity of ant antennae consist in the increase of apical segments of flagellum in relative size, so that the apical segment is the largest (by thickness, length), occasionally the last segments of flagellum make up for a club (see Fig. 1). In my opinion, this structure is explained by the same reasons: advanced spatial orientation (in the present case, due to an extension of the role of olfactory analyzers accompanied by a smaller contribution of vision unlike in flying hymenopterans), brood care, and subsequently communication via olfaction. These two selection vectors (increase in efficiency of orientation and control over perioral space) lead to the increase in number and density of olfactory receptors on apical segments of funiculus. The case is that olfactory receptors can be distributed in a hymenopteran flagellum almost uniformly throughout its length (although it is not a strict principle), however in ants they are accumulated both on apical and preapical segments of antennae, particularly clear this tendency is exhibited in myrmicines – their basal segments of funiculus are nearly devoid of olfactory sensilla (Hashimoto, 1990; Nakanishi, 2009; Euzébio 2013; Trible et al., 2017). The crucial importance of olfactory receptors in the ant phylogenetic branch, in comparison with other insects, including hymenopterans, is confirmed by the results of studies of the brain and genomes of ants (Guo and Kim, 2007; Gronenberg, 2008; Zhou et al., 2012). The number of genes responsible for olfactory receptors in ants exceeds several times, occasionally by an order of magnitude, those in other insects. The presence of 340-400 such genes has been shown for the studied ant species of different subfamilies and with different kinds of social organization (Dolichoderinae: *Linepithema humile*, Formicinae: *Camponotus floridanus*, Ponerinae: *Harpegnathos saltator*, Myrmicinae: *Pogonomyrmex barbatus*), besides the fruit fly *Drosophila melanogaster* has only 61 such genes, the honey bee *Apis melifera* ca. 170, and the parasitoid wasp *Nasonia vitripennis* ca. 300. It has also been shown that olfactory receptor genes in ants constitute a variable and rapidly evolving group of genes. According to experts, such values, particularly compared to other eusocial hymenopterans, indicate a more complex communication system in ants, based specifically on chemical reception. Experiments on gene shutdown with olfactory receptors have confirmed that workers which lack ability to smell demonstrate a dramatically reduced or completely lost ability not only to orient by smell, but also to communicate with relatives, to follow the pheromone trail, and care of larvae, although they are capable to feeding (Yan et al. 2017). Thus, the evolution aimed at the optimization of an organ, comprising olfactory receptors, raises no questions. Intensification of functions of spatial orientation in antennae of ants is confirmed indirectly by the relative reduction (decrease in the size) of optical analyzers (compound eyes), as well as by the brain structure – in modern ants, their olfactory processing zones are developed in a greater degree than visual (Gronenberg and Hölldobler, 1999; Gronenberg, 2008). Indeed, the closest flying relatives of ants have really enormous eyes; many stem ant taxa from Burmese amber also have relatively large eyes in comparison to modern ants, however they retain a non-ant structure of antennae, besides anteriorly shifted large bulgy eyes of modern ants are linked to a lifestyle of individual foraging (*Myrmoteras, Gigantiops, Myrmecia, Harpegnathos*). It should also be noted that, unlike some representatives of Dolichoderinae and Formicinae, ocelli are practically absent in workers of Myrmicinae and Ponerinae which are predisposed to live in the litter that indicates a tendency for the loss of ocelli in specialized litter dwellers. In this regard it is important to note that ants of different phylogenetic lines use absolutely different organs and substances to leave a pheromone trail: secretions of poisonous gland, Dufour gland, Pavan gland, mandibular glands, special leg glands; hindgut excretions and so on (Morgan, 2009). Such diversity testifies to an independent and multiple occurrence of “trailing” behavior in the crown ant lines, which is based on olfactory differentiation and mechanisms of advanced orientation, according to the hypothesis proposed here. So, the advancement of antennae for orientation purposes have served an impulse to the enhancement of chemical communication, i.e. has become a morphological preadaptation for further evolution of ant sociality.

According to the study of cranio-mandibular systems (CMSs) of ants by Dlussky and Fedoseeva (1988), presence of at least three mandibular teeth, triangular shape of the mandibles, and well-developed mandibular muscles also assign to the features shared by crown ant groups; apparently, they appear to be basic modalities of mandibles in crown ants. The authors assumed that three-toothed mandibles better coped with tasks of both holding prey and manipulating objects, that is necessary for ants, since they carry their larvae, in contrast to other hymenopterans. These arguments seem well founded. In this regard, I suggest that the presence and morphology of basal mandibular teeth in ants with the most enormous mandibles (Haidomyrmecinae) indicates that the workers could use them for carrying their larvae (Fig. 1*a*, 3*b, c*). Similarly, primitive social contemporary ants *Harpegnathos* sp. use the basal tooth of their giant holding mandibles to carry the larvae (Fig. 3). Yet, it is significant that winged reproductive females evidently possess the same mandibles (Barden and Grimaldi, 2016), (i.e. polymorphism of worker caste is absent), that points at a primitive social organization. Thus, such application (carrying larvae) could have been occurred a while earlier than the efficient tool for this function, it corresponds to the perceptions of the mechanism of morphological evolution.

**Fig. 3.**
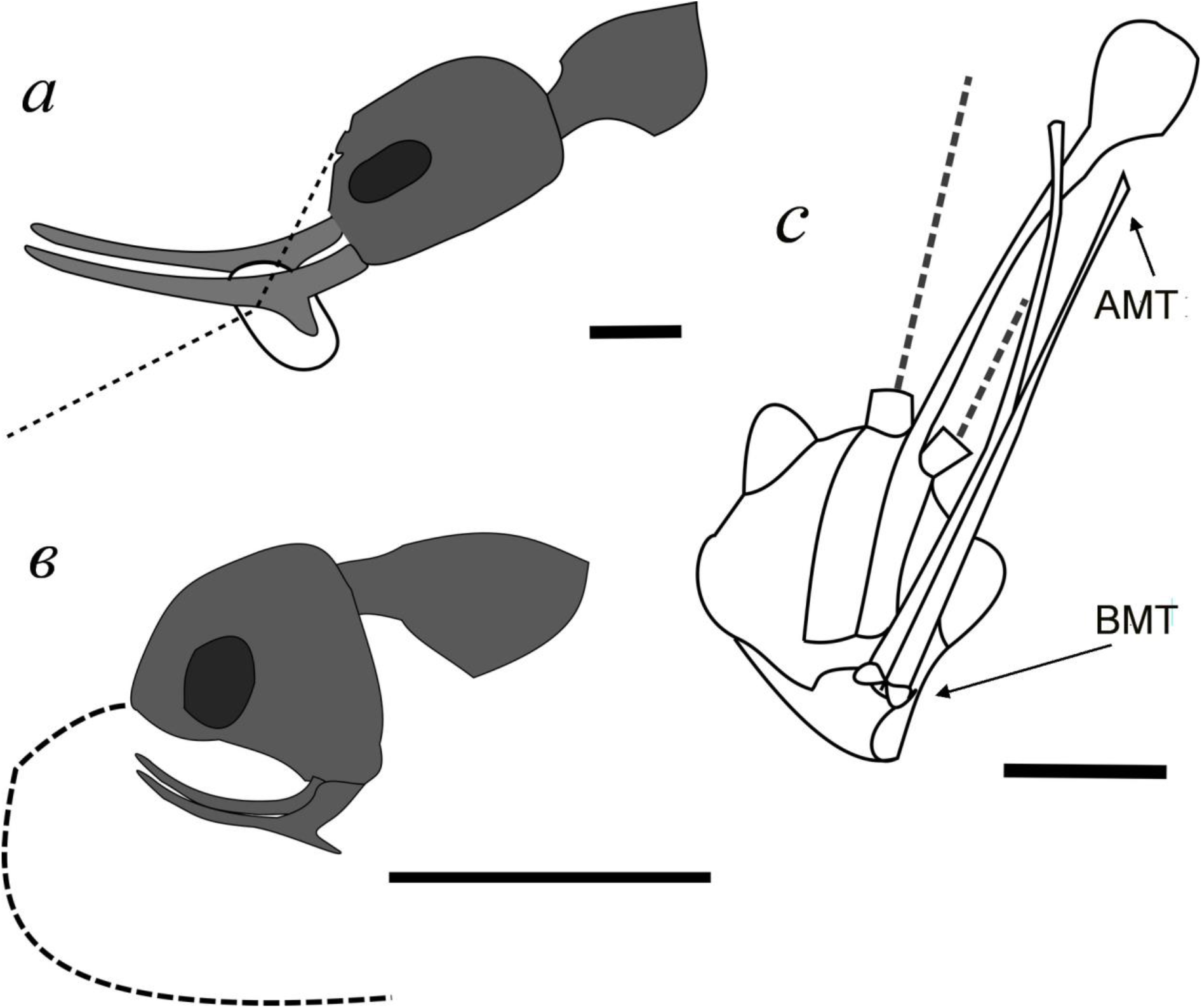
Mandible features of recent and Cretaceous extinct ants: *a* – application of basal teeth for egg and larva transportation in the long-mandible ant *Harpegnathos* sp.; *b* – mandible drawing of *Haidomyrmex zigrasi* Cretaceous ant in profile with a distinctive basal tooth position; *c* – head of *Ceratomyrmex ellenbergeri* (AntWeb: IGRBU002) from ventrolateral view. Abbreviations: AMT, BMT – apical and basal mandibular tooth. Scale bar: 1 mm.

The additional mandibular teeth exhibits in different hymenopteran lineages. For example, solitary leaf-cutting bees *Megachile* and mason bees *Chalicodoma* (Megachilidae), *Vespa* and *Vespula* (Vespidae) possess mandibles with several small teeth, that apparently is associated with nest-building behavior; the vespids can also use them for efficient holding and dissecting prey. The leptanilloid ant CMSs (mandibles with 3-5 teeth) look somewhat different, they better serve for holding prey rather than building a nest, therefore the question of the origin of multi-toothed mandibles in ant ancestors requires additional research. However, it appears that the invention of three(and more)toothed mandibles with developed musculature and special (compared to other insects) mandibular mobility (Richter, 2020) has become one of the crucial adaptations that allowed crown ants to occupy dominant positions in communities. Dlussky and Fedoseeva (1988) have shown that types of mandibles and possible pathways of evolution of their shape in ants can be correlated to the lifestyle of a species together with the presence of pronounced polyethism and polymorphism, besides the operating mechanism of the mandibles can be significantly modified. Thus, species with a primitive social organization are predatory with specialized mandibles (like holding or trap-type CMSs), then specialized individuals, soldiers, of nomadic ants possess hook-like smooth mandibles while ordinary workers have leptanilloid mandibles, and so on. Specifically both combination of a new shape of the mandibles and enlargement of the mandibular muscles indicate a better, more subtle control of movements, that fits well with the evolutionary picture of advancement of manipulations against the background of the above described transformations in anatomy and morphology of the antennae (approach to mandibles, density of olfactory receptors in apical segments). Ants are able to subtly manipulate even with very large trap-jaws. For example, long trap-jaw mandibles of *Odontomachus* sp. perform one of the fastest movements in the animal world when they snap shut catching and killing a prey, however, the workers are able to subtly manipulate with such very jaws for the purposes of careful accurate carrying and shifting eggs and early instar larvae in their nest (Just and Gronenberg, 1999). Particular significance together with pronounced vector of evolution, targeted at the advancement the manipulative capabilities of mandibles, are confirmed in crown ants by the results of Gronenberg’s studies (Gronenberg, 1996; Gronenberg et al. 1997; Just and Gronenberg W., 1999; Paul and Gronenberg, 2002 et al.). In a series of works he has shown the entire complexity of ant CMSs: changes in the length of muscles along with the ratio of different types of muscle fibers determine modifications of mandibles and also provide fine-tuning of functioning of these structures. He also has demonstrated that, in contrast to other hymenopterans, subesophageal ganglion of ants is considerably developed for the purposes of enhanced control of mandibular movements. Therefore, the structure of the mandibles (denticulate margin) and the presence (and development) of various mandibular muscles should be considered as one of significant factors for the formation of sociality in ants. Muscles affect the head shape, enhancing the action of the mandibles, even in the case of small mandibles (Fig. 4*b, c*). A different pattern is observed in stem ants (Fig. 4*d, e*).

**Fig. 4.**
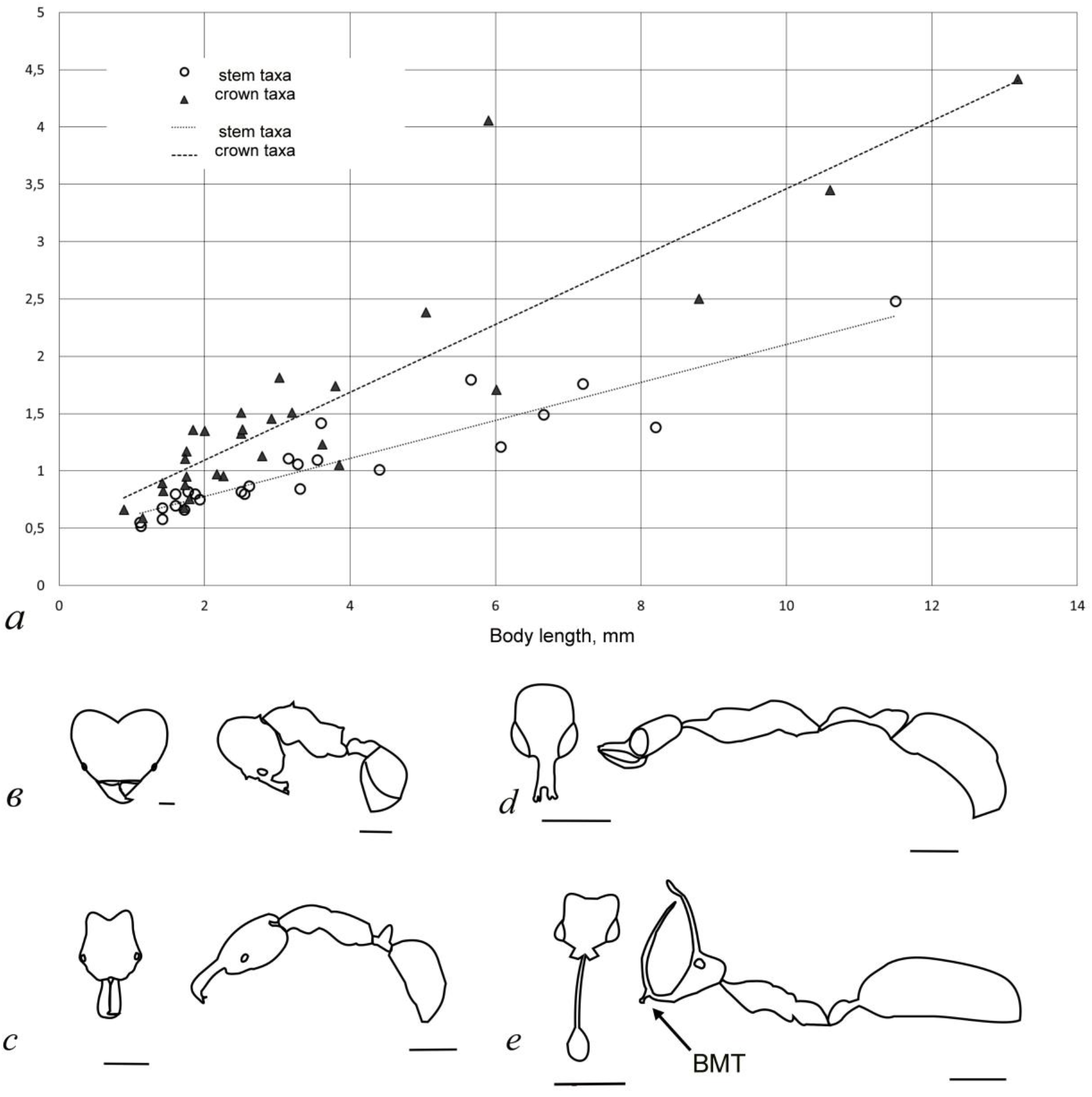
Size ratios and habitus features of recent and Cretaceous extinct ants: *a* – dependence between body length and head length (y-axis) in stem and recent ants; *b - d* – head dorsal and body profile contours: *b* – *Atta laevigata* (CASENT0922055); *c* – *Anochetus* sp. (CASENT0010781); *d* – *Aquilomyrmex huangi* (NIGP171999); *e* – *Ceratomyrmex ellenbergeri* (NIGP164022, NIGP164022). Scale bar: 1 mm. Abbreviations: BMT – basal mandibular tooth.

The unique diversity of formicoid mouthparts from Burmese amber, as has been repeatedly shown, exceeds the bounds of contemporary morphospace of recent ants, it stirs interest not only from the view of morphology, but also in regard to ecological (ethological, biocenotic) prerequisites for their occurrence (Barden and Grimaldi, 2016; Barden at al., 2020; Cao et al., 2020; Lattke and Melo, 2020; et al.). The design of the head capsule of Haidomyrmecinae representatives together with their remarkable curved mandibles indicate a different operating mechanism of their CMS (see Fig. 1*a*, 3*c*, 4*d, e*) in contrast to modern ants. Barden and Perrichot with colleagues, on the basis of external morphology and the unique finding of *Ceratomyrmex ellenbergeri* with a captured prey, a nymph of *Caputoraptor elegans*, have come to conclusion that mandibles of these representatives «*uniquely articulating in a vertical plane oblique to longitudinal axis of body, in addition to a moderate lateral opening»* (Barden at al., 2020). Recently discovered and yet undescribed species Colotrechninae sp., a representative of Chalcidoidea, exhibits a striking external resemblance to Cretaceous Haidomyrmecinae in head structures (facial processes, elongated setae near the mouthparts), mandibular morphology (elongated, curved upwards) and their relative position (Van de Kamp et al., 2022). The study of the only specimen has revealed the presence of a single, specifically anterior, mandibular condyle that allows mandibles to articulate in different planes – vertical movements (holding objects between the head capsule and mandibles), also lateral movements (manipulations in the space between mandibles). All studied members of the superfamily also have a unicondylar joint. The authors of the study associate such design of mandibular joint with evolutionary plasticity and striking species diversity of the group Chalcidoidea. Morphology of chalcid mandibles, specifically the presence of an additional tooth at the top of mandibles along with the developed denticulate margin at their basal part, as well as lifestyle of chalcids point at their superficial similarity to Cretaceous Haidomyrmecinae. However, the study of both the biology of this new insect and the mechanism of its mandibular movements, together with the comparative analysis of both modes of mandibular articulates and CMSs operation in extinct and recent ants, promise good prospects for uncovering biological peculiarities of Haidomyrmecinae. Also, Lattke and Melo noted similarity of head shape of Haidomyrmecinae with some parasitoid apocritan hymenopterans (*Tyrannoscelio, Stentorceps, Nanocthulhu*) (Lattke and Melo, 2020). Besides recently found Neotropical *Tyrannoscelios* possess long strong mandibles with several teeth that can articulate in a vertical plane (and slightly in an inclined plane), presumably for digging the ground. These facts point at the possibility of such rearrangements of mouthparts in hymenopterans. Most probably that the articulation of mandibles of Haidomyrmecinae executed not strictly in a vertical or a horizontal plane, but ventrolaterally, since it is necessary to spread mandibles to sides in order to use their basal teeth; this can also be evidenced by apical teeth diverged from each other in species with most long mandibles (see Fig. 3*c*) in order not to let mandibular tips interfere with one other when opening the jaws. Moreover, some peculiarities of mandibles of stem ant taxa provide evidence that this articulation characterizes not only Haidomyrmecinae, but also representatives of Zigrasimeciinae and Sphecomyrminae, at least *Gerontoformica*. It appears to me that predisposition to evolve in this direction lies in several morphological features of stem ants, specifically two-toothed mandibles and a weak, imperfect CMS. Two-toothed mandibles of hymenopterans fix the prey, preventing its rotation around the axis at the stinging. This is enough for flying predators, hunting even for large prey – after being stung the prey is motionless, it is fixed during transportation by air (such behavior can be observed in *Ammophila*, for example). But a flightless social predator needs to transport prey to the nest over substrate, and hence it can be robbed by competitors. The solution implemented by sphecomyrmines and the other stem taxa consists in pressing the prey to their head capsule (frons, clypeus), in contrast to recent ants which clamp objects between the mandibles. In Cretaceous formicoids, an increase in the size of prey affects lengthening of the apical tooth, enlargement and armouring of frontal space – the prey appears placed specifically between mandibles and head capsule. As the apical tooth of mandibles increases, the basal tooth remains closer to the base of mandibles (and to mouth opening). In some cases the basal tooth is exposed to some modifications. Evidently the basal tooth was applied to more subtle manipulations – probably manipulations with brood, dissection of prey in the nest or building the nest (see Figs. 1*a*, 3*b, c*, 4*e*). Thus, large mandibles are necessary in a greater extent not for killing, but as a means to fix the prey for stinging and further transporting to the nest. Modern ants have strong mandibles and the corresponding strong muscles, that is reflected in size and shape of their head capsule (see Fig. 4*b, c*); the second advantage of modern ants is group foraging, when tiny individuals with small jaws are able to protect and transport the prey regardless of its size (Fig. 5). The mandibles of hell ants, in spite of their length, most commonly do not look strong, furthermore the shape of their head capsule does not reflect the increase in muscle volume (like in modern ants – their size of head capsule remains relatively small (see Fig. 3) unlike processes (horns)). The fragility of their mandibles has also been noted by Lattke and Melo (Lattke, Melo, 2020), in contrast to thick, rigid mandibles of *Tyrannoscelio*. The “mandible conception” proposed here suggests that the serrated mandibular margin together with bristles and hairs on head processes are necessary for prey fixation and its control. Besides, setae and serration are also present in long-horned and long-jawed haidomyrmecines (*Ceratomyrmex*), in representatives of medium-sized sphecomyrmines with small mandibles (*Gerontoformica*), and in the smallest representatives of the stem groups with tiny mandibles (*Zigrasimecia*) (see Fig. 1*b, d*). Unlike stem ants where the increase in size of strictly two-toothed mandibles is determined by elongation of the apical tooth only, mandibles of modern ants increase in size throughout the entire length and the teeth are also distributed along the entire length, resulting in becoming strong structures (see Fig. *1a, g, m, n*; Fig. 3*c*; Fig. 4*e*).

**Fig. 5.**
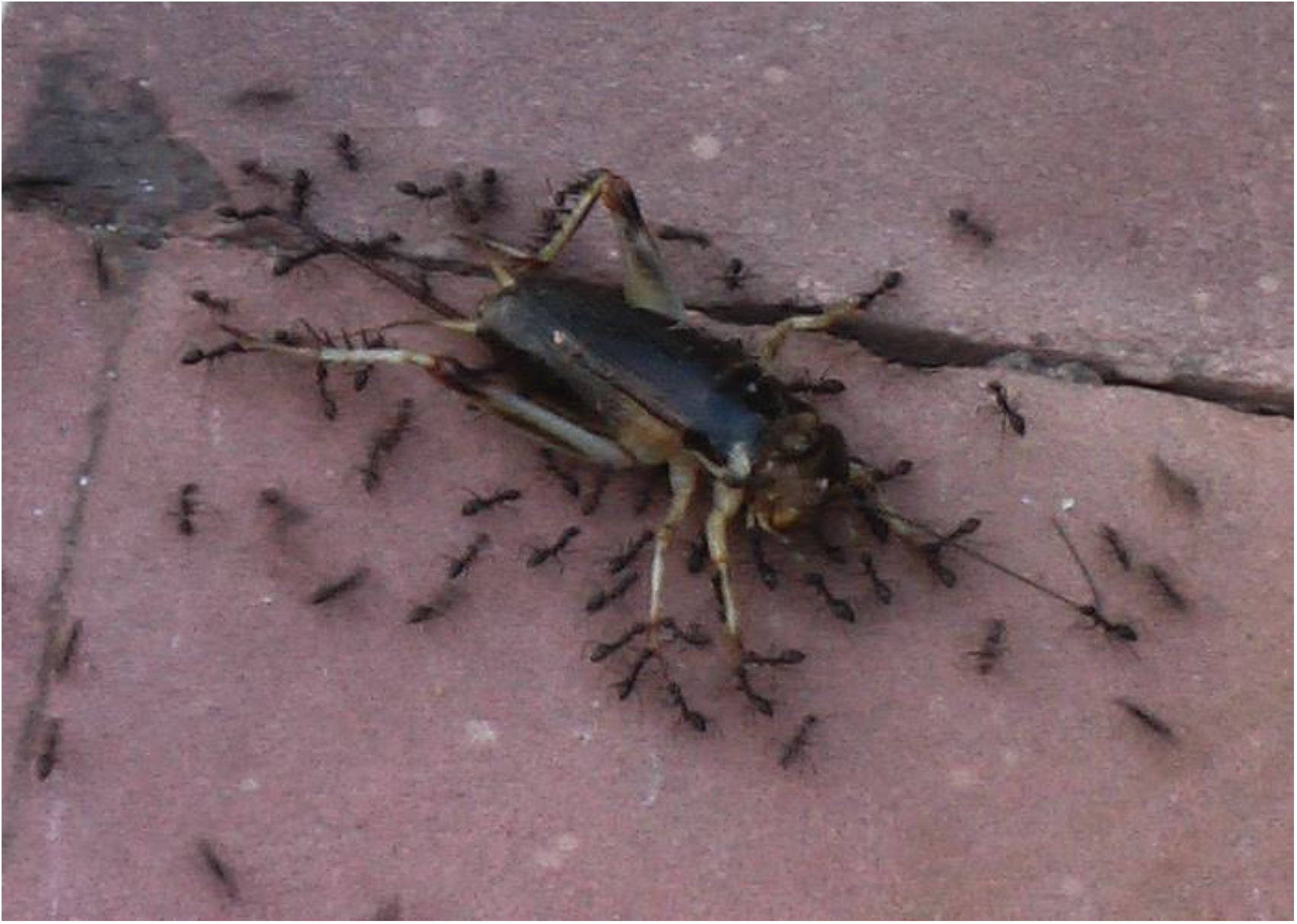
Group foraging and cooperative transport of large prey in small recent ants comprise several stages: prey detection, ant mobilization from nest to prey, prey killing, protection of prey, actual transportation, and partition of prey.

On the basis of the provided arguments, I consider the “trap-jaw” explanation, expressed yet at the time of description of the first representative of Haidomyrmecinae (*H. cerberus*) (Dlussky, 1996), imperfect, since the presence of strong muscles is needed for it, particularly in the regard of possible striking a prey. A “hemolymph feeding” hypothesis seems to be inconclusive. First, long and especially curved jaws are not excellent for hemolymph consumption (see Fig. 1*a*, 3*b*), it suffices to pierce a prey or bite it through with a short sharp “tool” (for this purpose, modern ants successfully use mandibles of any shape (*Mystrium camillae, Adetomyrma venatrix, Amblyopone silvestrii* etc.). Secondly, it is quite difficult to imagine the removing a prey off such “sabers” and their subsequent cleaning. The third argument is that feeding on liquid food implies its transmitting to other colony members, that is quite difficult to do with such jaws, also it could not be characteristic for socially primitive stem ant taxa in the framework of the conception proposed here. It should be also considered that feeding on liquid food implies its transportation in large amounts in the abdomen, which should be stretchable, therefore it would be interesting to estimate this parameter in the inclusions. More plausible in my opinion is the application of relatively short and thick mandibles for fixing and killing prey (including its piercing) followed by its dissection and consumption like in contemporary representatives of *Myrmoteras*, for example (multiple piercing, consumption of a grinded substance) (Moffett, 1986). Feeding on liquid food, such as honeydew or plant exudates, probably could be common among the stem taxa as being an additional nutrition, like in some poneromorphs which do not use oral trophallaxis, but feed “themselves on the road” or use (weakly effective) technique of droplet transportation between the mandibles (Paul and Roces, 2003; Novgorodova, personal communication).

The design of unique mouthparts of zigrasimeciines can be considered in the same view: setae, spicules and chitinous brushes evidence not of a unique way of hunting (or prey) of these ants, but just a mode of holding small soft-bodied invertebrates, which apparently could make up the diet of these tiny ants of body length ca. 2 mm. The authors of the descriptions of *Zigrasimecia hoelldobleri* and *Protozigrasimecia chauli* consider small invertebrates to be their most probable prey (Cao et al. 2020), too. Some authors suggest that both presence of a sting and a pronounced sculpture of a head capsule in such small insects evidence of rather predation than consumption of liquid food.

Crown taxa have taken a different path – manipulating objects between the mandibles only, excluding the head capsule involvement, and succeeded, by varying size, shape, and number of teeth of mandibles, on the one hand, muscle volume and muscle groups of the CMS, on the other.

The occurrence of an acid gland with the ability of spraying acid in some crown groups, according to Wilson and Hölldobler, is associated with a diet change (prey types?), but the relationship of this ability with the diet (change of prey types) has not been characterized (Wilson and Hölldobler, 2005). It appears that the ability of spraying acid can be claimed only in the case of coordinated collective actions, while an individual forager is more efficient having a sting. Although Smith’s remarkable observation has shown the advantages of acid attack of *Formica archboldi* against the powerful strong individual forager *Odontomachus brunneus* with trap-jaws (Smith, 2019), even so the acid attack appears a better defensive strategy in order to protect food resources and the nest in highly social ants: both as a poisonous agent and an alarm signal (Iakovlev, 2010). Possibly the development of both an acid gland and an ability of spraying secretion is linked to the sociality in a greater extent than to the diet, it is supported rather by the fact that specifically highly social members of the Myrmicinae, such as *Crematogaster*, have acquired this ability, than the subfamilies Dolichoderinae and Formicinae.

### Burmese amber orictocenosis as a model of the process of origin of modern myrmecocomplexes. Gibson’s principle

It is certain that the biodiversity of Burmite paleobiocenoses is very high. It is possible to state that modern biocenoses of this territory appear to be direct “descendants” of the Cretaceous ones. However, the taxonomic composition of Burmite ants and in similar contemporary ecosystems of this region differs at the level of subfamilies. What could be the reasons for the change in the taxonomic composition? The replacement of the gymnosperm flora was undoubtedly an important factor, but on its own it does not explain the dramatic changes of stem taxa to crown taxa – in what way glaciation cycles and thermal optima (together with changes in the floristic composition of biocenoses) in the Cenozoic led just to a change in the dominating genera, but not to a change in morphological “organization” of Formicidae. To explain the hypothesis proposed here, I have resorted to the insertion of the “Gibson’s principle” in the quality of the brief formulation of a tendency, well-known in the field of morphological evolution, evolution of biocenoses, and evolution of sociality (Treanore et al., 2021) (most probably, also the common feature of evolution at all its organization levels: molecular evolution, ontogeny evolution, macroevolution), to form something new not “instantly” and not “gradually”, but in a mosaic way, in the course of a search through combinations of elementary units of “organization”. These evolutionary phenomena can be perfectly described by the famous saying of an American-Canadian speculative fiction writer William Gibson: *“The future is already here – it’s just not evenly distributed.”* On the basis of Gibson’s principle, I propose a hypothesis of origin and formation of the modern biocoenotic role of ants, under conditions of food (ecological) specialization of stem ant taxa and occurrence of a set of crucial adaptations of crown ant taxa, by the example of Myanmar paleocenoses. Being an active, diverse group, stem ant taxa “formed” an adaptive space divided into ecological subniches, forbidden to other active arthropod predators. It can be assumed, in accordance with Gibson’s principle, that stem ant taxa exhibited the process of “formicoidization” over the Cretaceous: various characteristics of morphological and social organization in different combinations. However, a set of crucial adaptations – possibility of group foraging, effective communication, coordination of actions at the colony level, morphology of the CMS and antennae – has displayed to the full extent specifically in crown ant groups in the result of interrelated morphological changes and behavioral patterns; this has offered the possibility to the latter of becoming generalists and at the same time more efficient predators than ants of stem taxa. All of these things have provided competitive advantages over specialist ants of stem groups. Indeed, food specialization is present among modern ants: a) in some taxa with a primitive social organization, with small colonies (where tendencies of increase of relative mandibles size often persist, often the very same dwellers of the litter and upper layers of the soil – myrmicines and ponerines); b) in specific environmental conditions with sharply limited layering and resources, for example deserts (absolute size differentiation in workers among species); c) in much later formed in tropical forests with mortmass predominance, specifically in mushroom growers. Dominant in contemporary ecosystems ant taxa are presented by the species with large colonies (developed social organization and communication system), a wide food range which includes the keeping of insects that excrete honeydew (developed forms of trophallaxis) and collective predation non-specialized in prey type (Fig. 5). Most probably, the ants with a primitive social organization can afford none of the above listed points. Both the habitus of an effective fast terrestrial flightless herpetobiont (dendrobiont) predator and the social structure facilitated the autocatalytic process of improvement of a new life form. According to Dlussky, apparently at that time the radiation of the crown taxa proceeded, i.e. the adaptation of the new form to different strata of biocenosis accompanied by the replacement of the stem taxa. In spite of specialization, ants of stem taxa cannot hold their positions, because, as has been shown, advanced social skills contribute to more efficient control of food resources (I’Anson Price et al., 2021). So, poneromorphs have not been displaced from the main positions, as the DSH suggests, they were always restricted to the same place (where they originated) as now, where the resource base keeps the existence of not large colonies of predators (occasionally specialized) with individual foraging and a small proportion of carbohydrates in the diet – geobionts, stratobionts, and herpetobionts in a rich (forest) warm-climate biocenosis.

Judging by logical explanation of the background and morphological analysis, our hypothesis amounts to the following. Representatives of the stem ant taxa possessed a less developed communication ability, a less efficient CMS, and rather visual spatial than olfactory orientation, that prevented the development of sociality relying on odor stimuli (the development of communication ability, an increase in colony size, polyethism and polymorphism). Because of primitive communication organization their ecological role of predators among arthropods, in conditions of abundance and diversity of resources, resulted in an adaptive radiation via food specialization, that is a common evolutionary trend in insects (in particular, solitary hymenopterans), particularly in modern ants with a primitive social organization. Food specialization, in its turn, on the background of not flying, but cursorial, life form with a poorly developed communicative ability and a specific two-toothed CMS, though lacking a developed caste structure of the colony, determines an adaptive morphological evolution of species: relative size and variety of mandibles, size of individuals. A set of crucial adaptations of the crown groups – possibility of complex communication and coordination of actions, morphological peculiarities of the CMS and antennae – occurres due to reinforcing of the role of olfactory analyzers for the needs of a cursorial social herpetobiont insect, well-orienting in three-dimensional space, and results for its possessors in the formation of efficient non-specialized predators, given the possibility to control over the resource base as well as use and allocate within the colony (owing to trophallaxis) a new resource – liquid food (honeydew of Hemiptera, nectar). In the adaptive space organized by stem ant taxa the ecological niches, divided by specialists according to size and type of prey, are “formatted” in a new way by representatives of crown groups – and result in a hierarchical system, considered by modern ants as dominance (dominant, subdominants, influents), where type and size of prey are determined by the size of a colony, not the size of mandibles (Zakharov, 1991, 1994, 2015). However, ant mandibles can be modified to serve specific tasks even within the colony (polymorphism: workers and soldiers) due to advanced CMS and polyethism (polymorphism), under conditions of efficient allocation of food and separation of functions within the colony.

Thus, it seems that the change in myrmecofauna from stem to modern taxa happened to be on the score of occurrence of a new progressive group of ants, not owing to a diet change (due to the consequences of the replacement of gymnosperms by angiosperm plants), as suggested in the DSH. The diet change (consumption of liquid food, in particular), on the other hand, occurred due to the morphological and ethological evolution of the crown taxa. Since in line with Gibson’s principle, the environment was preadapted for the formation of a set of features embodied in crown ants due to mosaic distribution of features and characteristics among specialized stem ant taxa, which appeared less effective in this space and could not withstand competition.

## Acknowledgements

I express thanks to Antropov A.V. and Fedoseeva E.B. (Zoological Museum of Lomonosov Moscow State University), Chaika S.Yu. (Lomonosov MSU, Dept. of Entomology), Grinkov V.G. (Lomonosov MSU, Dept. of Biological evolution) for help in discussion of some details of morphology and providing with literature on hymenopterans.

Translated into English by Belyaev O.A. (Lomonosov MSU, Dept. of Entomology).

## Compliance with ethical standards

The article does not concern any research using warm-blooded animals as study objects.

## Footnotes

1 At the time of writing the article (2005) ponerine ants were considered as all subfamilies of the Poneromorph complex, which was eventually recognized as polyphyletic and divided into several subfamilies, i.e. Amblyoponinae, Ectatomminae, Ponerinae and other. Yet, Wilson E. supposed and considered the probable polyphyletic nature of the taxon, he also offered exactly ecological interpretation of the group, apart from taxonomic (cladistic) interpretation as insignificant.

